# Autonomic arousals contribute to brain fluid pulsations during sleep

**DOI:** 10.1101/2021.05.04.442672

**Authors:** Dante Picchioni, Pinar S. Özbay, Hendrik Mandelkow, Jacco A. de Zwart, Yicun Wang, Peter van Gelderen, Jeff H. Duyn

## Abstract

During sleep, slow waves of neuro-electrical activity engulf the human brain and aid in the consolidation of memories. Recent research suggests that these slow waves may also promote brain health by facilitating the removal of metabolic waste, possibly by orchestrating the pulsatile flow of cerebro-spinal fluid (CSF) through local neural control over vascular tone. To investigate the role of slow waves in the generation of CSF pulsations, we analyzed functional MRI data obtained across the full sleep-wake cycle and during a respiratory task during wakefulness. This revealed a novel generating mechanism that relies on the autonomic regulation of cerebral vascular tone without requiring slow electrocortical activity or even sleep. Therefore, the role of CSF pulsations in brain waste clearance may, in part, depend on proper autoregulatory control of cerebral blood flow.

**One-Sentence Summary:** Autonomic regulation of cerebral vascular tone mediates CSF flow and may contribute to brain waste clearance.

## Introduction

The clearance of metabolic waste products from the brain may be an important factor in maintaining brain health. While not well understood, the process by which this occurs is increasingly thought to involve the glymphatic pathway [1] and to be particularly active during sleep [2]. A recent fMRI study proposes a novel mechanism by which the brain itself may orchestrate this clearance. During sleep, electrocortical slow waves were found to be followed by joint cerebral-blood-volume and cerebro-spinal fluid (CSF) pulsations that may support clearance through the glymphatic pathway [3]. The mechanistic explanation for this sequence of events suggests that cerebral blood volume changes result from an active hemodynamic response to electrocortical activity, a key contributor to the fMRI signal [4, 5]. Slow waves are associated with a brief (200-500 ms long) but widespread reduction in electrocortical activity also known as a “down state” [6], which is accompanied by a reduction in total cerebral blood volume which in turn leads to the pulsatile inflow of CSF in order preserve the constant volume of fluids in the head [7](**Fig. 1**, “neural pathway”). These newly discovered CSF pulsations resulting from an active vascular mechanism are apparently neurogenic and possibly larger than those originating from the passive vascular response to blood pressure variations associated with cardiac and respiratory cycles. The latter also have been proposed to drive brain waste clearance [8–10].

**Fig. 1.**
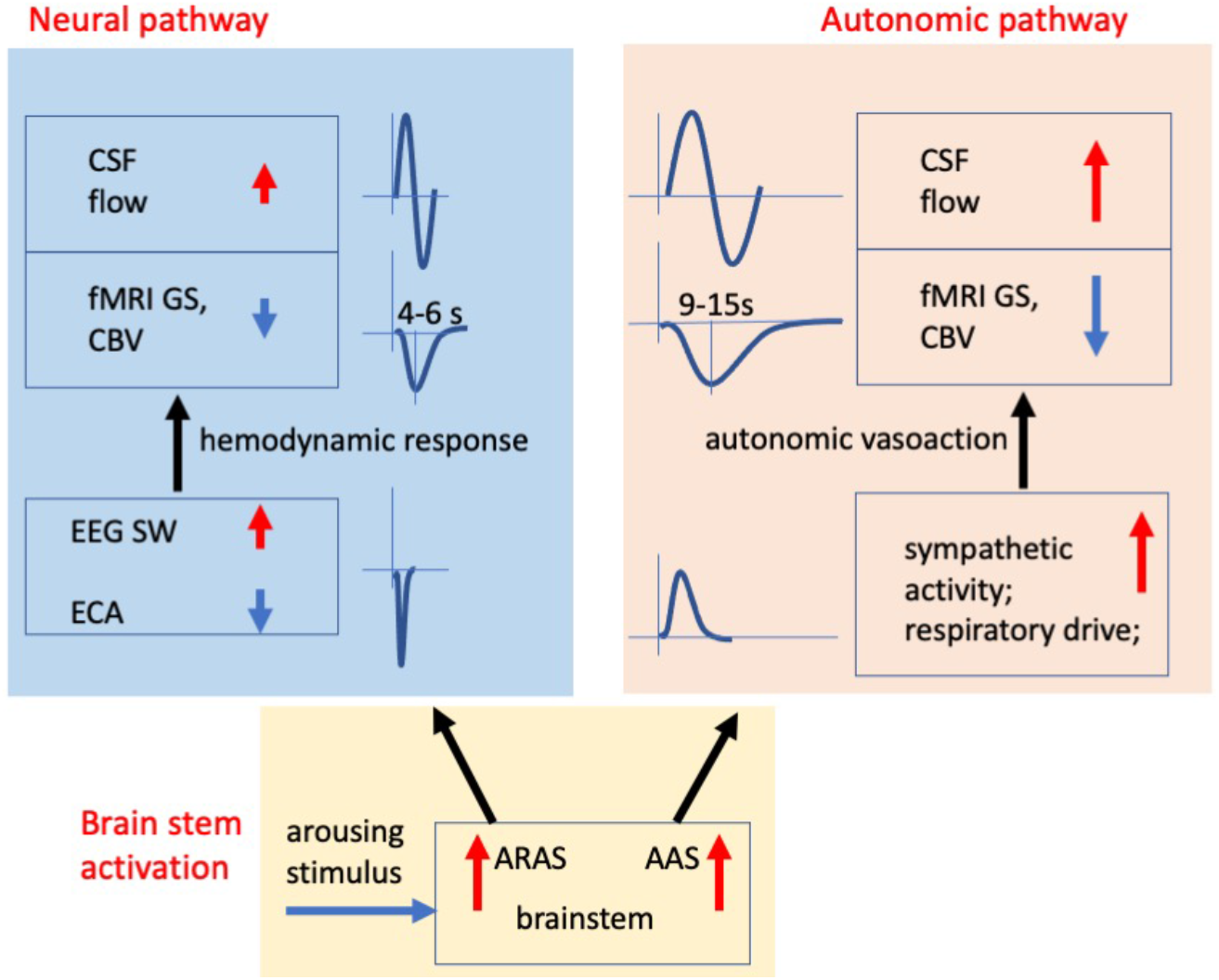
Mechanistic model for the generation of vasoactive CSF pulsations. CSF pulsations may be generated from intermittent vasoconstriction either elicited by widespread, episodic drops in electrocortical activity (ECA) (“*neural pathway”*), for example during an EEG slow wave, or through brief changes in autoregulation (“*autonomic pathway”*). In the neural pathway, vasoconstrictions lag reductions in electrocortical activity by the well-established 4-6 s delay dictated by the hemodynamic response [11, 12]. Autonomic pathway delays are longer and may reach 12-15 s, owing to the more sluggish effects of sympathetic and respiratory activity on vascular tone [13, 14]. During NREM sleep, joint changes in ECA and autonomic activity may be orchestrated by joint activations of (cortical) arousal and autonomic arousal promoting regions in the brain stem (“*brain stem activation*”), whose substrates partially overlap [15]. This may, for example, be in response to an arousing stimulus, either from the external environment, or internally generated (e.g. gastric stimulus, autonomic/baroreceptor signal, etc.). AAS = Autonomic Arousal System (brain stem neural substrate supporting autonomic arousal); ARAS = Ascending Reticular Activating System (brain stem neural substrate supporting cortical arousal). GS = global signal, CBV = cerebral blood volume, EEG = Electro-Encephalography, SWA = slow wave activity.

This generative pathway would expand the established role of slow waves in synaptic homeostasis and memory consolidation [16, 17] also to include orchestrating the clearance of waste products. This explanation would be consistent with the observation that the rate of fluorescent tracer clearance depends on the prevalence of slow waves in anesthetized mice [2, 18], and with the hypothesized relationship between Alzheimer’s disease pathology and changes in slow wave sleep (i.e. “deep”, or stage N3 non-REM sleep, the sleep stage with highest level of slow wave activity; see **SM Note 1**) in man [9, 19].

Nevertheless, the necessity of slow waves or even sleep for these novel, vasoactive CSF pulsations (and any associated clearance) has not yet been demonstrated, leaving room for alternative mechanistic explanations for brain waste clearance. For example, during sleep, major changes in autonomic physiology occur, including those associated with autonomic arousals [20, 21] (**SM Note 2**). Like neural activity, these can affect cerebral blood volume (and by inference CSF flow) in a widespread manner through vasoactive mechanisms not requiring slow waves or electrocortical activity, including CO_2_-mediated vasodilation and sympathetic vasoconstriction [4, 22] (**Fig. 1**, “autonomic pathway”). Autonomic arousals are relatively frequent during stages N1 and N2 (“light”) sleep (**SM Note 1**), the condition most prevalent during the original discovery of vasoactive CSF pulsations [3]. In fact, brief changes in heart rate, sympathetic activity, and inspiratory depth are known to co-occur with K-complexes [23–25], the type of slow wave that defines N2 sleep. These furthermore have been tied to seconds-long, widespread reductions in fMRI signal (and by inference cerebral blood volume) [26]. Could it be that the previously reported association between slow waves [3] and CSF pulsations is the result of brief changes in autonomic activity, for example autonomic arousals triggered jointly with K-complexes by activation of the brain stem (**Fig. 1**, “brain stem activation”)? If so, this may imply that an additional, autonomic mechanism exists for the generation of CSF pulsations through vasoaction, and possibly that not a lack of sleep, but rather compromised autonomic regulation of cerebral vascular tone affects brain waste clearance through CSF pulsations. To investigate this, we compared fMRI data with measures of autonomic physiology obtained across the full sleep-wake cycle and during a waking autonomic challenge.

## Results

To explore the association between slow waves, autonomic events, and CSF pulsations, we first re-analyzed data from a previous overnight sleep study, which included Blood Oxygenation Level Dependent (BOLD) fMRI, Electro-encephalography (EEG), as well as autonomic indicators. The latter included respiratory flow rate (RFR) as derived from a sensor measuring chest circumference (“chest belt”), as well as peripheral vascular volume (PVV) derived from finger skin photoplethysmography, an indicator of sympathetic vasoconstrictive activity [27]. A total of 59.5 hours’ worth of data from 12 subjects, distributed across all sleep states, was included in the analysis (**Table 1**, see **Online Methods** for data selection). In our analysis, the fMRI global signal (GS) was used as a proxy for total cerebral blood volume (**SM Note 3**), whereas the fMRI signal in the fourth ventricle and its connecting cerebral aqueduct revealed CSF flow into the brain (while being insensitive to flow out of the brain, see (**SM Data Analysis**). Spectral filtering in the 0.33-2.0 Hz band was used to extract slow wave activity (SWA) from the EEG signal across all sleep stages (**Online Methods**).

**Table 1:** Distribution of analyzed data over arousal states from 12 subjects in the sleep study

|  | N3 | N2 | N1 | W | REM |
| --- | --- | --- | --- | --- | --- |
| # 5 min segments | 94 | 345 | 140 | 74 | 61 |
| duration (min) | 470 | 1725 | 700 | 370 | 305 |
| # subjects | 10 | 12 | 12 | 12 | 10 |

### N2 slow waves occur jointly with CSF pulsations and autonomic arousals

Initial inspection of the data indeed suggested that the joint occurrence of K-complex type slow waves, GS changes, and autonomic arousals reported previously during stage N2 sleep [26] was also associated with CSF inflow events (**Fig. 2**). Comprehensive (n=12) lag-dependent correlation analysis (**Figs. 3a,b**) indicates that CSF inflow trailed SWA by 6-10 s, while fMRI GS changes trailed SWA by 8-15 s. We also observed a strong negative correlation near zero lag between the derivative of fMRI GS and the CSF signal (**Fig. 3c**), which supports the mechanistic interpretation that cerebral-blood-volume decreases cause CSF inflow [3]. Significant correlation between SWA and CSF/GS was also seen outside N2 (**Figs. 4, S1**), and, for all arousal states except REM, the variance in fMRI signals explained by SWA reduced after autonomic correction based on the estimated relationship between the autonomic signals (RFR and PVV) and the fMRI signal (**SM, Online Methods**)(**Fig. 4**). For GS, these reductions were 36±32%, 68±14%, 70±15% for N3, N2, N1, respectively, while for CSF, they were 34±26%, 71±15%, and 50±36%, respectively (mean ± sd, Student’s paired t-test: p<0.05). This confirms the potential role of autonomic activity in the observed SWA-fMRI correlations. This confirms the potential role of autonomic activity in the observed SWA-fMRI correlations.

**Fig. 2.**
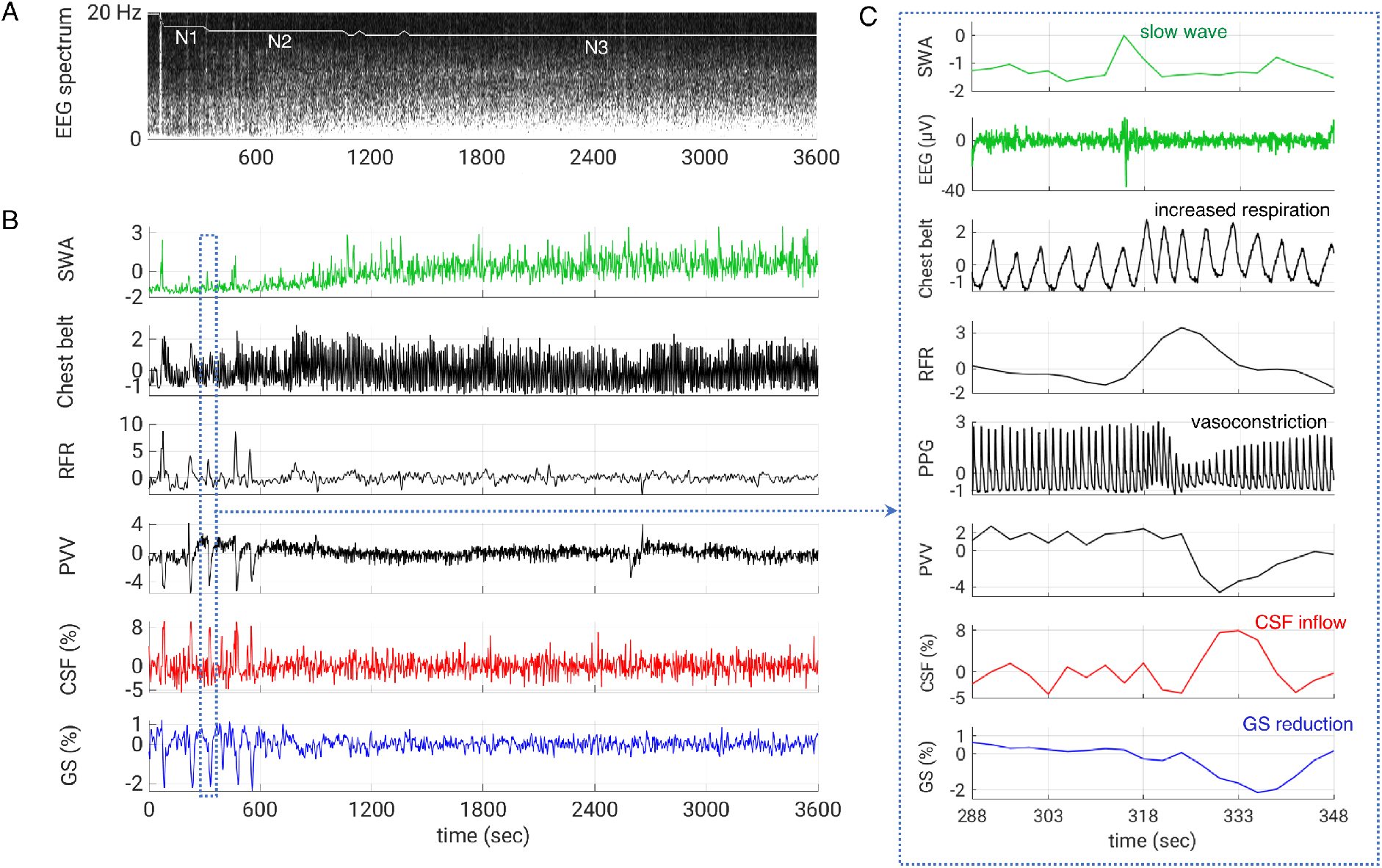
Association between slow wave activity (SWA), autonomic activity, fMRI global signal (GS), and CSF inflow signal during a one-hour segment of NREM sleep. (**A**) EEG spectrogram with identification of sleep stages. (**B**) During light (N1, N2) sleep, intermittent SWA attributed to K-complexes is followed by autonomic arousal, as evidence by increases in respiratory flow rate (RFR), a reduction in peripheral vascular volume (PVV), CSF inflow, and GS reduction. This association appears reduced during deep (N3) sleep, despite an increase in SWA. Signals are normalized to temporal standard deviation, except GS and CSF (% of mean). Inset (**C**) shows an expanded 60 s section to better appreciate the temporal relationship between the signals. PPG=raw photoplethysmography signal used to derive PVV.

**Fig. 3.**
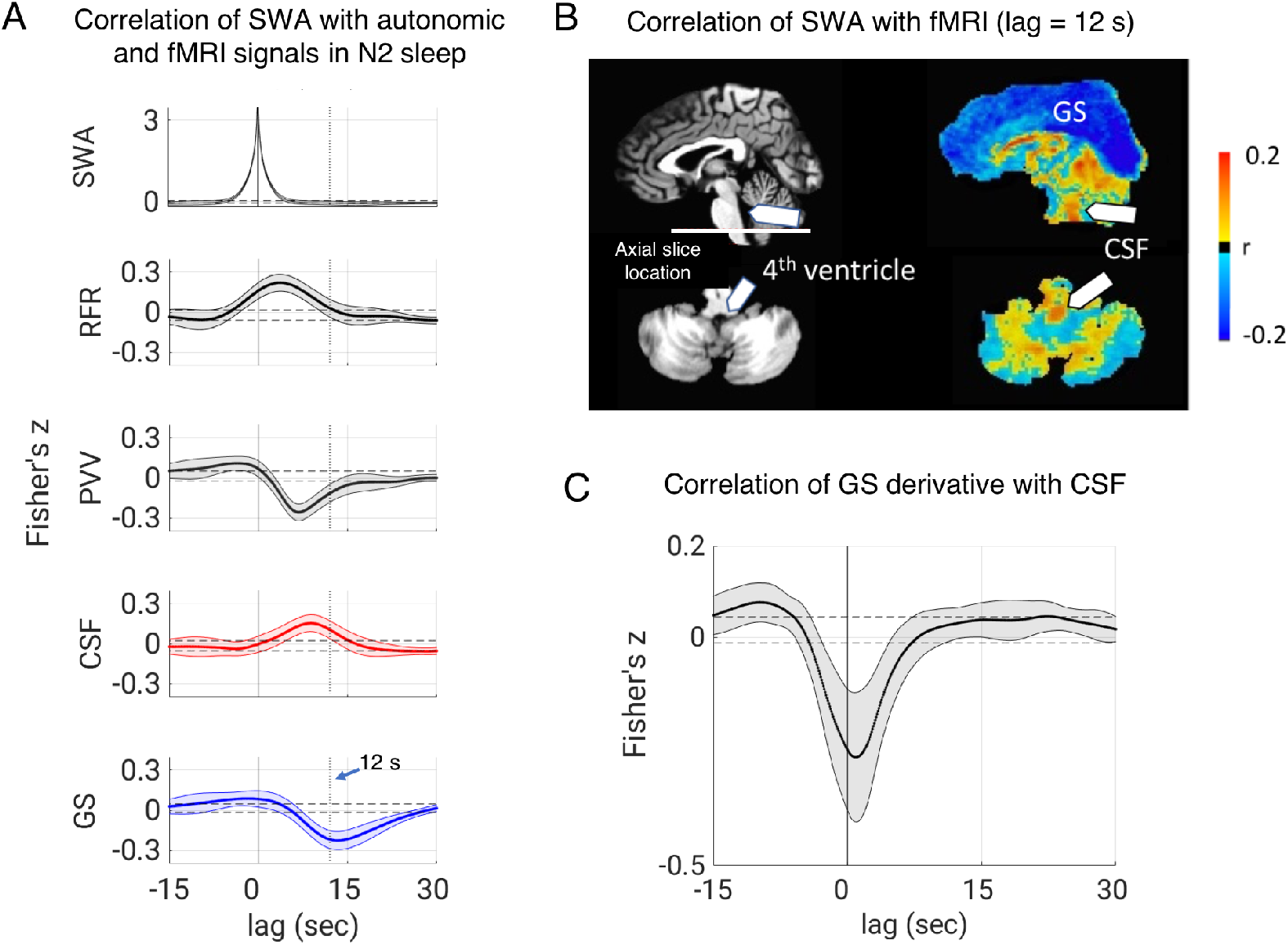
Association between SWA, autonomic activity, GS, and CSF during N2 sleep (n=12). (**A**) Lag-dependent correlation of SWA with autonomic and fMRI signals during N2 sleep (n=12). fMRI signals trail autonomic signals, which in turn trail SWA. (**B**) At a correlation lag of 12 s, SWA is associated with strong, widespread decreases in forebrain gray matter signal, while increases are observed in the fourth ventricle and cerebral aqueduct (arrows), indicating CSF inflow into the cranium. (**C**) Correlogram of the derivative of GS with CSF signal shows strong negative correlation near zero lag, consistent with cerebral blood volume reductions causing CSF inflow (**see Note 7**). RFR = respiratory flow rate, PVV = peripheral vascular volume, GS = global signals. Thick limes and shadings in A and C indicate mean and standard deviation across subjects respectively, while horizontal dashed lines mark the significance threshold (p=0.05).

**Fig. 4.**
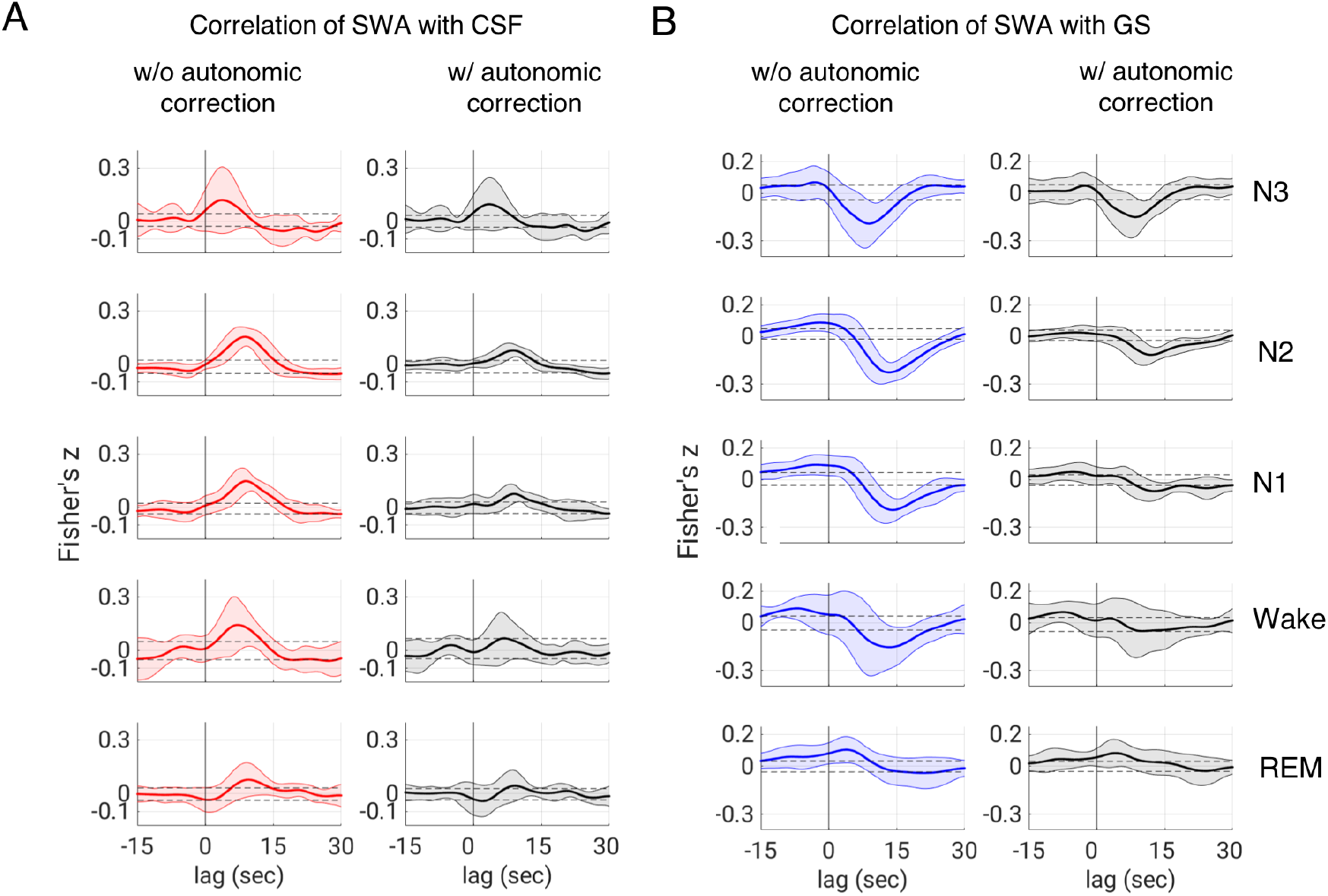
Sleep stage dependent correlation between SWA and fMRI. For both CSF (**A**, left column) and for the fMRI global signal (GS) (**B**, left column), significant correlation with SWA is observed for all arousal states except REM. Correction of the fMRI for autonomic signals (RFR and PVV) led to a reduction in correlation (see main text). Thick lines and shadings indicate mean ± sd across subjects. Horizontal dashed lines mark the significance threshold (p=0.05).

### CSF pulsatile activity does not follow the slow wave density increase with increasing sleep depth

While the correlation analysis presented above suggests that both autonomic events (i.e. brief changes in autonomic activity) and electrocortical activity may contribute to CSF pulsations, it does not report on their relative strengths across the various sleep stages or whether they possibly explain the previously reported association between the prevalence of slow waves (also called slow wave density, or slow wave power) and waste clearance [2, 18]. To allow for a more quantitative comparison, we estimated the fluctuation levels of CSF and GS from their temporal standard deviation and compared it to slow wave density (equal to SWA prior to detrending, see **Online Methods**). We found that going from N1 and N2 on one hand, to N3 on the other, slow wave density increased by 224±123% (mean ± sd, t-test: p<4*10^−6^) while in contrast CSF and GS fluctuation levels *decreased* by 14±12%, (t-test: p<8*10^−4^) and 19±22% (t-test: p<8*10^−3^) respectively. This suggests that SWA itself is not the dominant source of the observed CSF pulsations across sleep and that the previously reported increased brain waste clearance during deep sleep-like conditions in rodents [2, 18], if present in human, does not rely on CSF pulsations.

### Temporal sequence of events point to an autonomic contribution to CSF pulsations during NREM sleep

An interesting feature in our correlation data (**Figs. 3,4 and S1**) is the sizable lag of peak correlation between SWA and fMRI, as this suggests that an autonomic rather than neuronal mechanism is responsible for the CSF pulsations (see **Fig. 1**). To investigate this more closely, we quantified the lags of peak correlation across sleep stages. This required a somewhat stringent data selection, to avoid large timing errors due to poorly defined correlation peaks. For this reason, we only considered N1-N3 sleep, selecting subjects with more than 15 minutes of data and (absolute) correlations in excess of 0.1. For the SWA-GS correlation, this resulted in lags of 9.2±2.4 s (n=6, mean±sd), 13.1±2.8 s (n=12), and 13.6±2.8s (n=11) for N3, N2, and N1 respectively. Of note, the lag for N3 was significantly shorter than the lags for either N2 or N1 (p < 0.01 for both). These numbers were 4.8±1.5 s (n=6), 9.1±1.9 s (n=12), and 8.9±3.6 s (n=10) for the SWA-CSF correlation, with again the lag for N3 being significantly shorter than the lags for either N2 or N1 (p < 0.002, and p < 0.04 respectively).

This difference in lag-dependence between sleep stages was also apparent in results from spatially resolved correlation analysis, showing an early contribution (predominantly in posterior brain) at a lag of 6 s that appeared most pronounced during N3 (**Fig. S1**). Based on the well-established temporal characteristics of the hemodynamic response to brief electrocortical activity, one would expect the effect of a slow wave on the fMRI signal through the neurovascular response to peak after a delay in the range of 4-6 s [11, 12]. In contrast, the well-recognized fMRI autonomic contributions to the fMRI signal [28] are expected to peak at longer delays of between 9-15 s [13, 14]. Thus, the SWA-GS correlation lags in excess of 12 s found for N2 and N1 sleep are consistent with a dominant autonomic contribution, while the shorter lag for N3 (8.9±2.2 s averaged over the brain) may signify a joint contribution from neural and autonomic pathways (**Fig. 1**).

### Autonomic events alone can create CSF pulsations

While CSF pulsations secondary to vasoaction (either through a neural or autonomic pathway) may not explain the previously reported deep sleep dependent waste clearance, it remains possible that they play an important role outside deep sleep. In fact, the clearance attributed to CSF pulsations mediated by the passive vascular response to blood pressure variations with the cardiac cycle [29] or any such clearance related to the respiratory cycle would not be specific to deep sleep or sleep in general, and this may be also the case for vasoactive pulsations arising from challenges to the autonomic system. For example, independent of slow waves, respiratory challenges that alter intravascular CO_2_ will lead to cerebral-blood-volume changes [30, 31], and thus affect CSF flow. To investigate this possibility, we had alert subjects (n=11) take cued deep breaths, a task previously shown to lead to strong GS changes [13, 32]. Indeed, increases in respiratory flow rate were accompanied by spatio-temporal patterns of fMRI signal changes similar to those seen after slow waves during N1 and N2 sleep (**Figs. 5, S2**). The amplitude change of GS after inspiration averaged - 1.9±0.7%, consistent with previous work [13] when taking into account differences in experimental conditions (i.e. echo time and MRI field strength, see **SM Note 4**). Peak GS reductions following a deep breath occurred at correlation lags in the range of 13.2-14.8 s (mean 13.9±0.6 s), similar to the range of around 10-15 s reported previously [13, 14]. These results indicate that vasoactive CSF pulsations are not specific to sleep and may simply result from autonomic events, either through changes in RFR, or through sympathetic vasoconstriction.

**Fig. 5.**
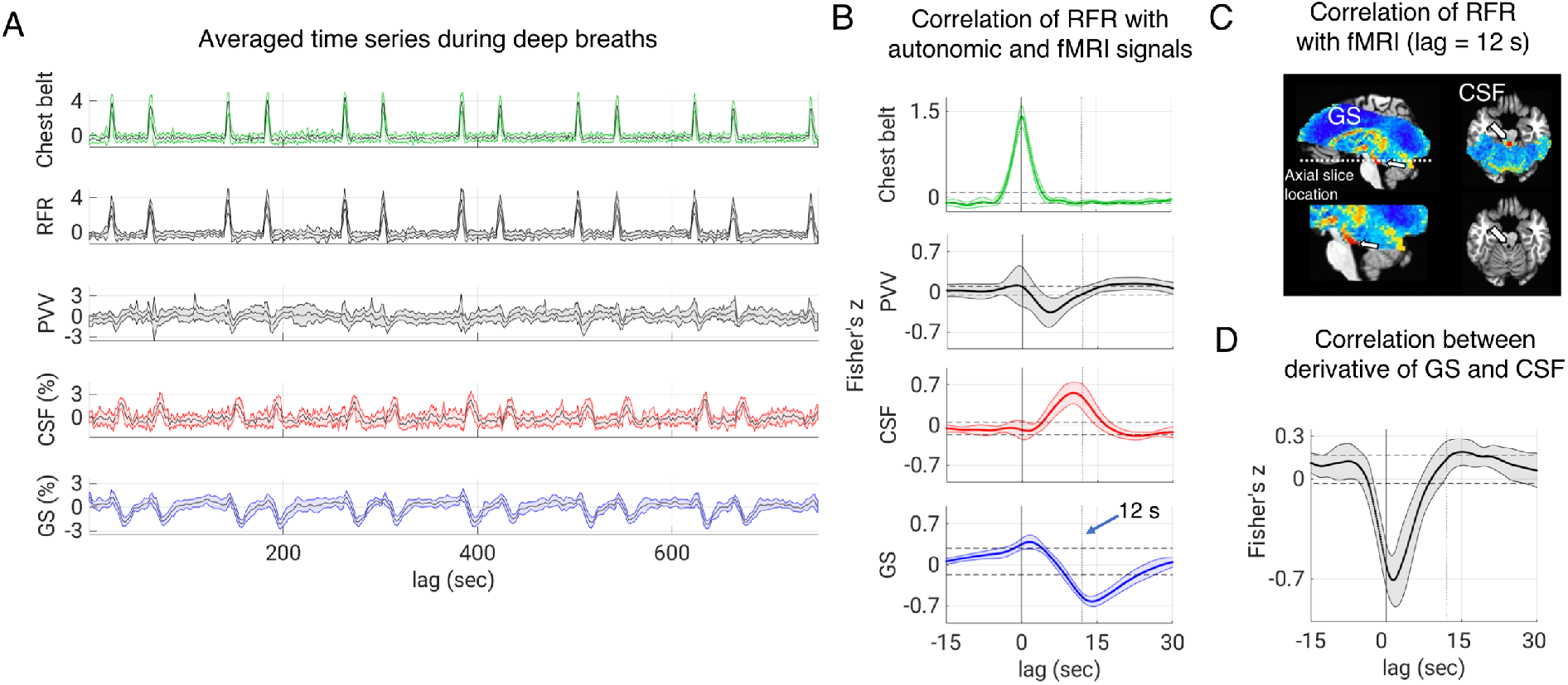
CSF pulsations in response to an autonomic challenge. Voluntary deep inspirations are associated with CSF signal increase and global signal (GS) decrease (**A**). Signals are normalized to their temporal standard deviation. (**B**) The correlation of respiratory flow rate (RFR) and fMRI signal mimics the results seen during N2 sleep (Fig. 3B), including a temporal precedence of autonomic changes relative to fMRI changes similar to those seen during N2 sleep. (**C**) At a lag of 12 s, an increase in RFR leads to a widespread reduction in fMRI signal. In contrast, CSF signal near the fourth ventricle shows a strong increase (arrows) due to inflow into the cranium. Correlograms of RFR with fMRI signals further confirm temporal precedence of autonomic changes relative to fMRI signal changes, similar to those seen during N2 sleep (Fig.3A). (**D**) As in Fig. 3C, a strong negative correlation between the temporal derivative of GS and CSF near lag 0 confirms the notion that the fMRI CSF signal reflects inflow resulting from a cerebral blood volume decrease. Thick limes and shadings in B and D indicate mean and standard deviation across subjects respectively, while horizontal dashed lines mark the significance threshold (p=0.05).

## Discussion

The correlation between brain waste clearance and slow wave density observed in anesthetized mice [2, 18] suggests that clearance in human may be particularly effective during N3 NREM sleep, when slow waves of electrocortical activity are most prevalent. A recent report, primarily based on analysis of fMRI data from the lighter (N1-N2) NREM sleep stages, suggested that this clearance may be mediated by CSF pulsations that directly result from SWA through a neuro-vascular response to electrocortical activity [3] (**Fig. 1**, “neural pathway”). Here we investigated whether these CSF pulsations indeed are preferentially present during N3 sleep, and to what extent autonomic mechanisms (**Fig. 1**, “autonomic pathway”) may contribute to their generation.

We revealed that CSF pulsations, while present during N3, may be stronger during the lighter sleep stages when slow waves are less prevalent. Furthermore, the amount of variance in the CSF flow signal explained by SWA reduced substantially after autonomic correction, particularly during N1 and N2 sleep, suggesting CSF pulsations do not require slow waves and in part may result from autonomic arousal or autonomic instability. This was confirmed with a respiratory challenge during wakefulness, which showed that deep inspirations are followed by spatiotemporal patterns of fMRI and CSF changes similar to those seen after slow waves during N1 and N2 NREM sleep. These findings that CSF pulsations secondary to active vascular mechanisms do not preferentially occur during N3 sleep, and do not require slow waves or even sleep, put into doubt the notion that they explain the previously observed N3 sleep-dependent waste clearance [2, 18].

The coincidence between autonomic arousals and slow waves during light sleep seen here is well established (for reviews see [23, 25, 33]) and can be explained in part as follows (**Fig. 1**). Slow waves during N1/N2 sleep present similarly widespread electrocortical activity as slow waves during N3, but generally originate from a different mechanism that involves the brain stem [6, 23, 33, 34]. For example, sensory input from environmental and internal (e.g. gastric) stimuli may simultaneously activate the ascending reticular activating system [35] and the autonomic arousal system [15, 36] because of their overlapping and highly connected neural substrates (see **Fig. 1** “brain stem activation”). The cortical response to this multi-faceted arousal is an EEG K-complex (also called “Type 1” slow wave [34]), whereas autonomic effectors are pupil dilation, CNS vasoconstriction, increased respiratory drive, and increased heart rate.

The relatively long lag between SWA and fMRI changes, in particular during light sleep, is suggestive of a dominant contribution from autonomic variability rather than variability in electrocortical activity. This can be understood by examining the duration of various phenomena. While the electrocortical activity reduction during a single slow wave lasts only 300-500 ms, autonomic arousals are more persistent and may last 5-15 s [37, 38]. As a result, the time-integrated effect of autonomic arousals on cerebral blood volume and CSF flow may be more substantial (see **SM Note 5**). During deep sleep, slow waves occur in more rapid succession (up to ~1/s), still the associated reduction in cortical activity may have only limited effect on CSF pulsations due to the sluggishness of the fMRI hemodynamic response (5-10 seconds width [11, 12]). Nevertheless, the shortening of the delay between SWA and fMRI going from N2 to N3 (see e.g. **Fig. 4**) suggests a non-negligible contribution of electrocortical activity on CSF pulsations. Thus, it appears there are (at least) two distinct vasoactive mechanisms responsible for CSF pulsations: one mediated through the neurovascular response to electrocortical activity, and one related to autonomic changes, mediated by sympathetic control of vascular tone and the vasodilatory effects of intravascular CO_2_.

Compared to previously reported CSF pulsations resulting from passive arterial and venous pressure effects from cardiac and respiratory cycles [8, 39], the newly discovered vasoactively generated CSF pulsations are of larger amplitude and therefore have the potential to be of more importance to waste clearance. Rough estimates of the volume of CSF (temporarily) displaced during each autonomic arousal can be obtained from previously reported CSF flow velocities in the 4^th^ ventricle as well as the cerebral-blood-volume and BOLD-fMRI changes previously observed with autonomic challenges. These estimates are in the range of 3-3.5 ml, which is substantially larger than the 0.5 ml and 1-2 ml estimates for the direct, passive effects on CSF flow from cardiac and respiratory cycle induced pulsations respectively (**SM note 6**). While the autonomic arousals (during sleep) and autonomic variability (during wake) are not as frequent as the rate of cardiac and respiratory cycles, it is possible that their larger associated CSF flow changes improves mixing of solutes in the CSF compartment, thereby facilitating waste clearance. To be sure, the above flow estimates are only a single aspect of the complex flow dynamics throughout the brain, and it remains unclear to what extent they reflect actual waste clearance.

Brain waste clearance through the glymphatic system is a multi-factorial process involving fluid influx into the brain, a mostly diffusional movement through the parenchymal interstitium [40, 41], and outflow in part along the arterial and venous vasculature [42–44]. The role of CSF pulsations in this process remains incompletely understood and a topic of ongoing research. Pulsations mediated through vascular volume changes may facilitate mixing of CSF solutes, beyond what occurs with diffusional and convective processes alone [2]. It is also possible that pulsations may result in a net flow of CSF and thus facilitate clearance of waste products [45]. Based on our findings, one would expect the largest CSF pulsations to occur with modulation of autonomic activity outside N3 (deep) sleep, when cerebral blood volume changes are largest and most frequent. The previously observed improved clearance during N3-like conditions may therefore not have resulted from increased CSF pulsations, but rather from other factors, for example an increase of convective fluid flow through tissue facilitated by the augmentation of extracellular space [2, 18].

A further possibility is that vasoactive CSF pulsations have relevance for brain waste clearance independent of sleep. Our results suggest that such CSF pulsations may be expected with the strong autonomic and autoregulatory modulations associated with e.g. postural changes, respiratory apneas, and arousals. Widespread neural activity changes with arousals may also contribute to the pulsations. Their CSF pulsation strength would, in part, depend on vascular reactivity to modulations in blood CO_2_ and sympathetic activity, while their frequency would, in part, depend on physical activity. Any pulsation-dependent waste clearance would then be diminished in individuals with compromised autoregulation or (neuro-)vascular reactivity, potentially exposing them to increased risk for pathology. Such a link may explain the previously reported findings of an association between cerebral amyloid angiopathy, a hallmark of Alzheimer’s disease, and compromised vascular reactivity [46–48]. If confirmed, this would provide an opportunity for use of the fMRI GS, and by inference vasoactive CSF pulsations, as a potential early biomarker of Alzheimer’s disease.

Future experimental studies, for example those using fluorescent tracers, may further clarify the interplay between hemodynamic autoregulation, the facilitatory role of CSF pulsations in waste clearance, and pathology.

Finally, it is possible that vascular pulsations themselves may contribute to brain waste clearance by facilitating solute drainage along vessels, independent of their potential role in promoting mixing or net flow in CSF spaces. Cardiac-cycle related pulsations for example may lead to transport of solutes along the basement membrane of smooth muscle cells in the arterial vasculature [29], and recent modeling work shows that slower types of pulsations may even be more effective [49]. Notably, a recent study in mice reported increased para-arterial clearance of a fluorescent dextran tracer with slow, spontaneous and task-evoked vascular activity [50].

## Acknowledgments

Susie Fulton, Steve Newman and, Jiaen Liu are acknowledged for advice and assistance. This work utilized the computational resources of the NIH HPC Biowulf cluster (http://hpc.nih.gov).

## Author contributions

Conceptualization: DP, PSO, JHD

Methodology: DP, PSO, JAdZ, PvG, HM, YW

Data acquisition: DP, PSO, JAdZ

Data analysis: DP, PSO, HM, JHD

Visualization: JHD, PO

Supervision: DP, JHD

Writing – original draft: DP, JHD

Writing – review & editing: DP, PSO, HM, JAdZ, YW, PvG, JHD

## Data and materials availability

The datasets analyzed for this study are available from the first or corresponding author upon reasonable request and in accordance with institutional policies.

## Online Methods

### Data acquisition

All data for this study were collected under human subjects research protocols that were approved by the local Institutional Review Board and involved obtaining informed consent from the participants. Data were collected in two different experiments (“sleep experiment” and “respiratory challenge”) that were not specifically designed for the questions of this study.

#### Sleep experiment

Data acquisition for a previously described sleep experiment [51] included two consecutive nights of concurrent fMRI/EEG data collection while subjects slept in a 3 T Siemens MRI scanner. For two weeks prior to the experiments, subjects had a regular sleep schedule as confirmed by home monitoring. No sleep deprivation protocols were applied. Whole brain (50 axial slices) fMRI data were collected at 2.5 mm spatial resolution (2.5 × 2.5 mm in-plane, with 2.0 mm slice thickness and 0.5 mm slice gap), 3 s temporal resolution, a 90° flip angle, and an echo time of 36 ms. Data were collected with multi-slice echo-planar imaging in a slice-interleaved fashion. EEG data was recorded at 5 kHz digitization rate from 64 channels covering most of the scalp using a Brain Products (Gilching, Germany) EEG system. Concurrently acquired peripheral physiological measures included chest belt (to monitor respiratory chest excursion) and finger skin photoplethysmography (PPG) (to monitor cardiac rate and peripheral vascular volume). These were acquired using a Biopac acquisition system with TSD200-MRI and TSD221-MRI transducers, and an MP 150 digitizer sampling at 1 kHz (Biopac, Goleta, CA, USA). Data collection for both EEG and peripheral physiology were synchronized with the fMRI through a trigger signal provided by the MRI scanner.

#### Respiratory challenge experiment

Autoregulatory effects on CSF pulsations were studied during a 12.5-minute fMRI experiment in which subjects (n=25) were visually instructed to take a brief deep breath at specific times (10, 50, 130, 170, etc., with successive 80 s and 40 s intervals) while being kept awake with a visual task. fMRI data (28-32 slices) covering part of the brain (see e.g. **Figs. 5, S2**) was collected on a Siemens 7 T MRI scanner at 2.0 mm spatial resolution, 2 s temporal resolution, flip angle 70°, and an echo time of 30 ms. As with the sleep experiment, chest belt signals and finger skin PPG were recorded as well using Biopac hardware.

### Data Analysis

#### Sleep experiment

All 12 subjects (out of a total of 16 attempts; age 18-35, 8 female) that completed both nights of scanning were analyzed. During each night, fMRI experiments were irregularly interrupted by acoustically stimulated or spontaneous awakenings. This resulted in a series of experimental runs that varied in length between 5 minutes and 3 hours. Only runs more than 30 minutes long and with good quality EEG, fMRI, and autonomic physiology data were analyzed, which constituted 5955 minutes’ worth of data. This was further reduced to 3570 minutes after excluding data with excessive motion and non-uniform or ambivalent arousal state (see below). The fMRI data were preprocessed using AFNI software (version 20.3.01)[52] and a tailored version of its “afni_proc” pipeline. It included the removal of outliers (spikes) and slow polynomial trends as well as two harmonics of the cardiac and respiratory signals from a modified RETROICOR model [53]; furthermore, slice-timing correction, motion registration, and non-linear alignment to the Talairach template (TT_N27_SSW) were performed. Finally, the estimated motion parameters and their derivatives were regressed out of the data while ignoring (censoring) any outliers of motion greater than an aggregate translation (mm) and rotation (degrees) parameter of 0.3 (see below for details on the censoring).

After preprocessing, further analysis was performed with IDL (Harris Geospatial, Broomfield, CO, USA). Correlation analysis was performed between EEG SWA (see below) on the one hand, and fMRI and peripheral physiological signals on the other. For fMRI signals, both single voxel time series as well as region of interest basis (ROI) averages were considered. The GS ROI covered the entire forebrain (both gray and white matter). This was done to capture blood volume changes in both tissue types, as their combined change is relevant for the CSF pulsations of interest. Alternatively, one could consider just the gray matter (where the strongest and most consistent effects are expected), however this would complicate the comparison between the sleep experiment and the respiratory challenge experiment due to differences in partial volume effects (because of their different spatial resolutions and geometric distortions). The CSF ROI included the area of the fourth ventricle (or its connections to the cerebral aqueduct and towards the spinal canal) in the bottom (5 most inferior) slices of the fMRI slice stack, which are most sensitive to inflow effects (see e.g. [3]). The slice at the very bottom of the stack was excluded because of its increased sensitivity to motion. The CSF ROI was defined for each subject on the fMRI data itself, in which CSF stood out because of its bright signal (as a result of its high proton density and long T_2_ relaxation time) (See example in **Fig. S3**). Substantial inflow sensitivity was expected because of the long T_1_ relaxation time of CSF (~ 4 s, weakly dependent on field strength, see [54]), which leads to substantial saturation of the magnetization under the conditions of both the sleep and the respiratory challenge experiments. As a result, inflowing (unsaturated) CSF will lead to a signal increase. For the respiratory challenge experiment (7 T, repetition time 2 s, flip angle 70°) saturation was estimated at 50%, while for the sleep experiment (3 T, repetition time 3 s, flip angle 90°), the estimate was 47% (remaining magnetizations of 50% and 53% respectively). As a result, inflowing (unsaturated) CSF will lead to an fMRI signal increase.

EEG signal was corrected for MRI gradient and cardio-ballistic artifacts and down-sampled to 250 Hz using Analyzer software (Brain Vision, Morrisville USA) as described previously [51]. Sleep scoring was performed from a central electrode in 30 s epochs according to standard criteria with standard filters and channel references [55]. Subsequently, SWA was calculated for successive 3 s time windows that matched the timing of the fMRI volumes. This was done by integrating the Fourier spectrum magnitude of the EEG time signal from a central electrode (C3, according to standard naming convention [56]), and then integrating the (magnitude) spectrum over the frequency range of 1/3-2 Hz [26]. This is the frequency range expected to contain the bulk of the spectral power associated with both K-complex and non K-complex type slow waves across all non-REM sleep stages [57]. Any outliers in the resulting SWA time course exceeding 4 standard deviations above the mean were set to the mean signal level.

Peripheral physiological measures were extracted as follows. First, to remove artifactual spikes from the data, time points exceeding 4 standard deviations above the mean were set to the mean signal level. Then, low-pass filtering was performed on the PPG and respiration belt signals at 30 and 10 Hz cut-off respectively. From the PPG signal, an indicator for peripheral vascular volume (PVV) was derived by calculating the standard deviation of the PPG signal in 3 s segments, which is a measure of the excursion amplitude caused by the cardiac beat and is proportional to the volume of blood detected by the PPG sensor [58]. The filtered chest belt signal was used to derive a measure of respiratory flow rate, which previously was found strongly to affect the fMRI signal throughout the brain [13]. This was done by taking the derivative of the low-pass filtered belt signal (see above), rectifying it, and applying a secondary low-pass filter at 0.13 Hz cut-off. Prior to this low-pass filtering, outlier data points in the derivative were suppressed by clipping the signal below 4 standard deviations. Importantly, these two signals derived from the PPG sensor and the chest belt should be considered as approximate indicators of peripheral vascular volume (PVV) and respiratory flow rate (RFR); they are roughly proportional to these autonomic processes but by no means quantitative.

To compare the relationships between fMRI, EEG, and autonomic measures across sleep stages, correlations were also performed on segments of fixed length (100 data points of 3 s each, duration 5 minutes). Data segments where sleep scoring was not possible due to residual MRI artifact or segments with excessive subject motion were excluded. The latter was based on the following analysis of the fMRI motion parameters (obtained from the motion correction procedure of the fMRI preprocessing): first, an aggregate measure of head motion was constructed by adding the standard deviation of all 6 motion parameters (3 translations in millimeters and 3 rotations in degrees). Then, frames with this aggregate (summed) standard deviation exceeding 0.5 units were excluded. This was done to reduce the effect of arousal state transitions on the analysis. An additional selection was made based on the uniformity of a sleep stage by excluding all segments with less than 90% of the time in a single sleep stage. Combined, these criteria led to the exclusion of 40% of the data frames; the distribution of remaining segments over sleep stages is reported in **Table 1**. To evaluate the contribution of autonomic processes on the correlation between SWA and fMRI signals, the analysis was repeated after regressing out the RFR and PVV signals. This was done for each segment separately using RFR and PVV at two time-shifted versions each. Shifts of 12 and 15 s were used for RFR and 0 and 3 s for PVV, values expected to be effective in accounting for physiological contributions to the fMRI signal [26].

Lagged correlation analysis was first performed voxel-by-voxel on data at the MRI temporal resolution (3 s for the sleep data, 2 s for the respiratory challenge data). Subsequently, results were selectively averaged within subjects according to dominant sleep stage. Finally, data were averaged across subjects, and the standard deviation was calculated across subjects, as well as estimated from permutation analysis, in which segments were shuffled within subjects. For the ROI-based analysis, signals were first interpolated to 0.1 s temporal resolution before correlation analysis to facilitate determination of lags at which peak (maximum or minimum) correlation occurred. Lags within the range of 0-20 s were used for this purpose.

#### Respiratory challenge experiment

From the 24 subjects that were part of this dataset, 15 consistently complied to the respiratory cues as judged from the respiratory belt signal excursions. This was based on their belt traces correlating with the respiratory paradigm with correction coefficients in excess of 0.7. Of these, 11 (mean age 26.8±4.7, 7 females) had sufficient fMRI brain coverage to include the fourth ventricle or the cerebral aqueduct and were analyzed for the relationship between the CSF signal, GS, and physiological measures. As with the sleep data analysis, slice timing, motion registration, and motion regression were performed with AFNI. CSF and GS calculation were performed analogously to the sleep analysis, with the exception that, due to the limited spatial coverage, the GS calculation sometimes did not include the most superior slices of the brain (for an example of the area covered by the fMRI slice stack, see **Figs. 5, S2**). In addition, the CSF ROI typically only included the CSF superior to the fourth ventricle inside the cerebral aqueduct.

Pre-processing of the peripheral physiological measures followed that of the sleep experiment with the exception that outlier detection was omitted to avoid interference with the relatively large signal excursions during the cued breaths. Inspiratory depth, as determined from the respiratory belt signal, was correlated with the various other signals, including the RFR, PVV, and the fMRI signal (MRI voxel-by-voxel data and MRI ROI data). This was done on the full data length of 375 time points (12.5 min). For the MRI voxel-by-voxel data, lagged correlation was performed at the fMRI temporal resolution of 2 s. For the ROI-based analysis, lagged correlation was performed after interpolating the data to 0.1 s temporal resolution. This facilitated peak timing analysis of the correlograms. Standard deviation was calculated across subjects.

### Statistical Tests

To test the statistical significance of the group level lagged cross-correlations, we estimated a 95% confidence interval based on percentiles (corrected for multiple comparisons via Bonferroni correction, number of comparisons = number of TRs corresponding to 45 s) of an empirical null distribution (**SM Figs. 3-5**). We created the null distribution by computing and pooling all cross-correlations from subjects. This is performed after converting Pearson’s r to Fisher’s z, and for each of the vector pairs by circular shifting but excluding lags of interest. For the sleep data, excluded lags correspond to the 45 sec intervals shown in the correlograms. For respiratory challenge data, which had a task paradigm with 40 s periodicity, we excluded 20 s data segments around the expected correlation peaks.

To estimate significance of subject level cross-correlations, we used the relation between Pearson’s r and t-scores, i.e., p-value is calculated using a t-distribution with n – 2 degrees of freedom. We marked the lags which give p < 0.05 (corrected for multiple comparisons, as described above) (**Fig. S2**). In addition, a paired Student’s t-test was used to compare correlations before and after autonomic correction at selected lags of interest (**Fig. 4**).

For the sleep stage specific analysis, results were first averaged within subjects, followed by analysis across subjects (n=12). For the data segments selected for analysis (**Table 1**), stages N3 and REM were only present in 10 of the 12 subjects.

## Supplementary Materials

### Note 1: Description of arousal states

Based primarily on EEG features, human arousal states can be classified according to established criteria [55]. By convention this is done in 30 s epochs. The stages include Wakefulness (W), Rapid Eye Movement sleep (REM, or R) and three stages of non-REM sleep (N1-N3). N1-N3 stages are associated with increasing sleep depth, with N1 and N2 typically considered “light” sleep and N3 “deep” sleep. The latter is also referred to as slow wave sleep. W is defined by a high frequency (> 7 Hz) EEG. W encompasses both eyes closed wakefulness with prominent 8-12 Hz waveforms as well as eyes open wakefulness with faster waveforms. REM is defined by three coincident phenomena: a) an EEG similar to N1, b) an EMG that is at the lowest level of the recording, and c) eye movements with an initial deflection ≤ 500 ms. During N1, 8-12 Hz activity in the EEG is replaced by 4-7 Hz activity. The addition of transient K-complexes/Type I slow waves (≤ 2 Hz, typically ≥ 75 μV) and/or Sleep Spindles (11-16 Hz) mark the beginning of N2. The first epoch of N2 is often considered unambiguous sleep onset. N3 is scored when ≤ 2 Hz, ≥ 75 μV waveforms occupy at least 20% of the 30 s scoring epoch. These waveforms are now called slow waves/Type II slow waves.

### Note 2: Definition of autonomic arousal

In scientific literature, the term “arousal” is used liberally and may have various meanings. In this report, in analogy with Halasz et al. [24], “arousal” in isolation and “cortical arousal” refer both to cortical arousal, which classically is defined as the switch from low frequency or “synchronized” EEG activity to high frequency “de-synchronized” EEG activity. “Autonomic arousal” refers to an episodic change in autonomic processes, as reflected in acceleration of the heart rate, sympathetic vasoconstriction, and increases in respiratory flow rate. “Brain stem arousal” refers to the activation of brain stem arousal centers that may lead to cortical and/or autonomic arousal. “Arousal state” refers to the various sleep stages (see **Note 1**).

### Note 3: fMRI as proxy for CBV

The BOLD fMRI signal results from an interplay between changes in cerebral blood volume (CBV) and changes in flow (and to a lesser extent changes in cerebral oxygen consumption). According to the influential balloon model [59], the time course of fMRI changes elicited by evoked neuronal activity closely (within 1-2 s) track that of the evoked CBV changes. This has been experimentally confirmed in both animal and human [60, 61].

### Note 4: Amplitude of fMRI global signal drop compared to previous work

The magnitude of the GS drop after a deep inspiration averaged 1.9±0.7% (TE=30 ms, 7 T). Previously, Birn et al. [13] found an average of around 0.8% (TE=30 ms, 3 T) for all voxels significantly responding to the respiratory challenge. Assuming a baseline T_2_* of ~50 ms at 3 T, 30 ms at 7 T, and a linear increase of inspiration-related R_2_* (=1/T_2_*) change with field strength [62] see also **Note 6**), our 1.9% effect size at 7 T translated into a DR_2_* of 0.63 /s, equivalent to a 0.27 /s DR_2_* at 3 T. The latter would lead to a ~0.8 % signal change at 3 T, consistent with that found by Birn et al. [13].

### Note 5: Visibility of the neuro-vascular effect from the change in electrocortical activity with slow waves compared to brief stimuli in event-related fMRI

This primary fMRI GS effect after a K-complex (i.e. slow wave during N1 and N2) had delays in the range of 10-15 s, while little effect was seen at the shorter lags of around 5 s typical of the hemodynamic response to neuronal activity. This may be somewhat surprising in light of the large EEG changes observed with K-complexes, and the substantial reduction in neuronal activity measured with invasive electrophysiology [6]. However, it should be realized that the duration of this neuronal activity change is only about 500 ms or less, which is relatively short compared to the neuronal activity changes evoked in typical task fMRI experiments (seconds long). In comparison, autonomic arousals typically lasted many seconds, which is at least an order of magnitude longer. This may explain the effect size difference of these two mechanisms. In addition, while neuronal firing may nearly completely cease during the negative deflection (i.e. “down state“) of the K-complex it may increase several-fold during stimulation tasks [5]. It seems therefore likely that the absolute change in the level of neuronal firing during a slow wave does not exceed that during a strong stimulus of equal duration.

### Note 6: CSF displacement from CBV changes from various sources

Any waste clearance mediated by CSF pulsations will depend on the frequency of the pulsations and the amount of CSF displaced. This is difficult to do based on inflow-mediated signal enhancement in fMRI as the latter relationship with flow velocity is rather complicated. To overcome this, one can estimate cerebral blood volume changes from the observed GS changes, and relate these directly to CSF displacement (see **Note 7**), as is done below.

#### Expected CSF displacement for the neuro-vascular effect from isolated slow waves

Based on prior work [31, 63], the estimated local CBV change with long (many seconds) strong stimulation (electrical forepaw stimulation in rat) ranges between 10-20%. Considering that the hemodynamic response is several seconds long (~5 s full width at half maximum), and assuming linear temporal summation of the hemodynamic response to neuronal activity [11, 64, 65], and a comparable magnitude change in neural activity during a K-complex and strong stimulation, one would expect the K-complex to lead to a several (> 3) fold smaller CBV change, or in the range of 3-6%. Assuming K-complexes affect all of human brain gray matter (an overestimate) occupying a total volume of around 800 ml in human, and about 4% (32 ml) of this gray matter being blood, one would expect a 1-2 ml maximum CBV decrease (and a negligible effect in white matter). This would correspond to an upper limit of 1-2 ml CSF influx, substantially smaller than that expected from autonomic mechanisms (below). A larger CSF influx may occur if the neuro-vascular mechanism also affects extra-parenchymal vessels, as these may contain a similar amount of blood as the intraparenchymal vessels.

#### Expected CSF displacement from a CO_2_ change with deep breaths

Autonomic changes may affect vascular tone through various mechanisms, including CO_2_ mediated local vasodilation and sympathetic vasoconstriction. Deep breaths reduce alveolar and intravascular CO_2_ and lead to vasoconstriction. Gray matter blood volume changes associated with inhaled CO_2_ gas mixtures may reach up to 45% (for a 5% CO_2_ or 35 mmHg pCO_2_ change [31, 66]). During a deep inspiration as performed in the current study, the expected momentary pCO_2_ change is (very roughly) 8-10 mm Hg [32]. This would correspond to a 10% CBV change and a 3 ml CSF influx, about double the estimated upper limit for the neurogenic change with a K-complex. Considering the fact that the gray matter involvement of the CO_2_ effect is likely more widespread than the gray matter involvement with the neural activity change associated with a slow wave, the CO_2_ effect may even be more dominant. Even larger differences between neuronal and autonomic mechanisms are possible when breathing changes co-occur with changes in sympathetic activity, as often is the case with K-complexes [67].

An alternative way to estimate the CSF displacement with a deep breath is from the observed BOLD changes based on the previously reported relationship between CBV and the fMRI signal [31]. The authors found (at 2.0 T) that a CBV change of +24% with electrical rat forepaw stimulation led to R_2_* changes of −0.54 /s, and these numbers were +17% and −1.1 /s at 4.7 T. Linearly interpolated to 3 T, this would mean a −0.8 /s R_2_* change for a +20% CBV change. Assuming a gray matter baseline T_2_* of ~50 ms at 3 T [62] this is equivalent to a signal change of 2.8% at TE=35 ms (sleep experiment). Linearly extrapolated to 7 T, with TE=30 ms (respiratory challenge experiment), assuming a baseline T_2_* of ~30 ms [62], this is equivalent to a 1.9 /s R_2_* change and a signal change of 5.9%.

In the respiratory challenge experiment, the observed whole brain (all tissues) signal change (averaged over breaths and subjects, standard deviation over subjects) was 1.9±0.7% (n=11). Considering the CBV change occurs predominantly in gray matter, and gray matter is only about a 60% fraction of total brain volume (i.e. white matter, gray matter, and CSF) [68, 69], global gray matter signal changes are estimated at about 3.5%. This then would correspond to a CBV change of 20%x3.5/5.9 = 12%, and a CSF inflow of about 3.5 ml (12% of 32 ml), in line with the numbers found above. As with the neurovascular effect, an even larger CSF influx may occur if CO_2_ changes also affect the blood volume in extra-parenchymal vessels, as these may contain a similar amount of blood as the intraparenchymal vessels.

#### Estimated CSF displacement from the observed GS change following K-complexes

The fMRI GS change following a K-complex varied greatly but typically reached an amplitude of about 1%. This is consistent with the ~ 2% change that was observed by Fultz et al. [3] (who also used TE≈30 ms and 3 T MRI), considering the difference in region used for GS calculation (here, the region included was about 40% white matter, where little effect is seen, whereas Fultz’s region focused on gray matter). Recalculating our GS change for just gray matter (assuming white matter has negligible effect) would lead to a signal change of 1.7%. Using the fMRI-CBV translation data from the previous section, this corresponds to a CBV change of 1.7*20%/2.8 = 12%, similar to that seen with the respiratory challenge experiment (above). The corresponding volume of CSF inflow would also be about 3.5 ml. This displacement is also within the range that can be calculated from the flow velocities in the fourth ventricle estimated in Fultz et al. [3], which were around 1 cm/s and occasionally higher. Considering the cross-sectional area of the fourth ventricle is around 1 cm^2^, this would correspond to a flow of about 1 cm^3^/s. With the flow response lasting several seconds (full width at half maximum response duration of > 5 s; see Fig. 4 in [3], or **Figs. 2b** and **3a** in this report), this would correspond to a CSF displacement in excess of 3 ml.

#### Estimated CSF displacement with K-complexes and deep breaths compared to that caused by blood pressure effects from cardiac and respiratory cycles

The vasoactive CSF pulsations described here and in Fultz et al. [3] may be substantially larger than those reported previously related to cardiac and respiratory cycles. In addition to CO_2_ mediated vasoactive effects of the intraparenchymal arterial vasculature, sympathetic effects may substantially contribute as well. Sympathetically mediated CNS vasoconstriction is thought primarily to affect the extra-parenchymal vasculature, whose total volume exceeds that of the intraparenchymal arterioles primarily affected by CO_2_. Joint occurrence of deep inspirations and sympathetic activity, for example during K-complexes, have synergistic vasoconstrictive effects and may lead to CSF displacements in excess of the 3-3.5 ml estimates based on deep inspirations alone. These effects are stronger than previously reported for the direct pressure effects of cardiac and respiratory cycles.

For pressure pulses related to the cardiac cycle, CSF movement has been estimated to involve a volume of about 0.5 ml [70]. During the inspiratory phase of the respiratory cycle, CSF flow velocities in the fourth ventricle may reach about 1 cm/s [71]. Again (see above), taking into account the cross-sectional area of the fourth ventricle (~1 cm^2^), and a 2 s long inspiratory phase (during only part of which maximal flow velocity is reached), total displaced volume may reach a maximum of about 1.5 ml [72]. This assumes maximal flow velocities are reached throughout the cross-sectional area.

### Note 7: The temporal GS-CSF relationship

As postulated previously [7], one expects the inflow of CSF into the cranium to correspond to the CBV change. Assuming constant morphology of the relevant CSF fluid canals, one therefore would expect the CSF flow velocity to be proportional to the derivative of CBV, here estimated by taking the derivative of the fMRI GS [3].

---

**Fig S1.**
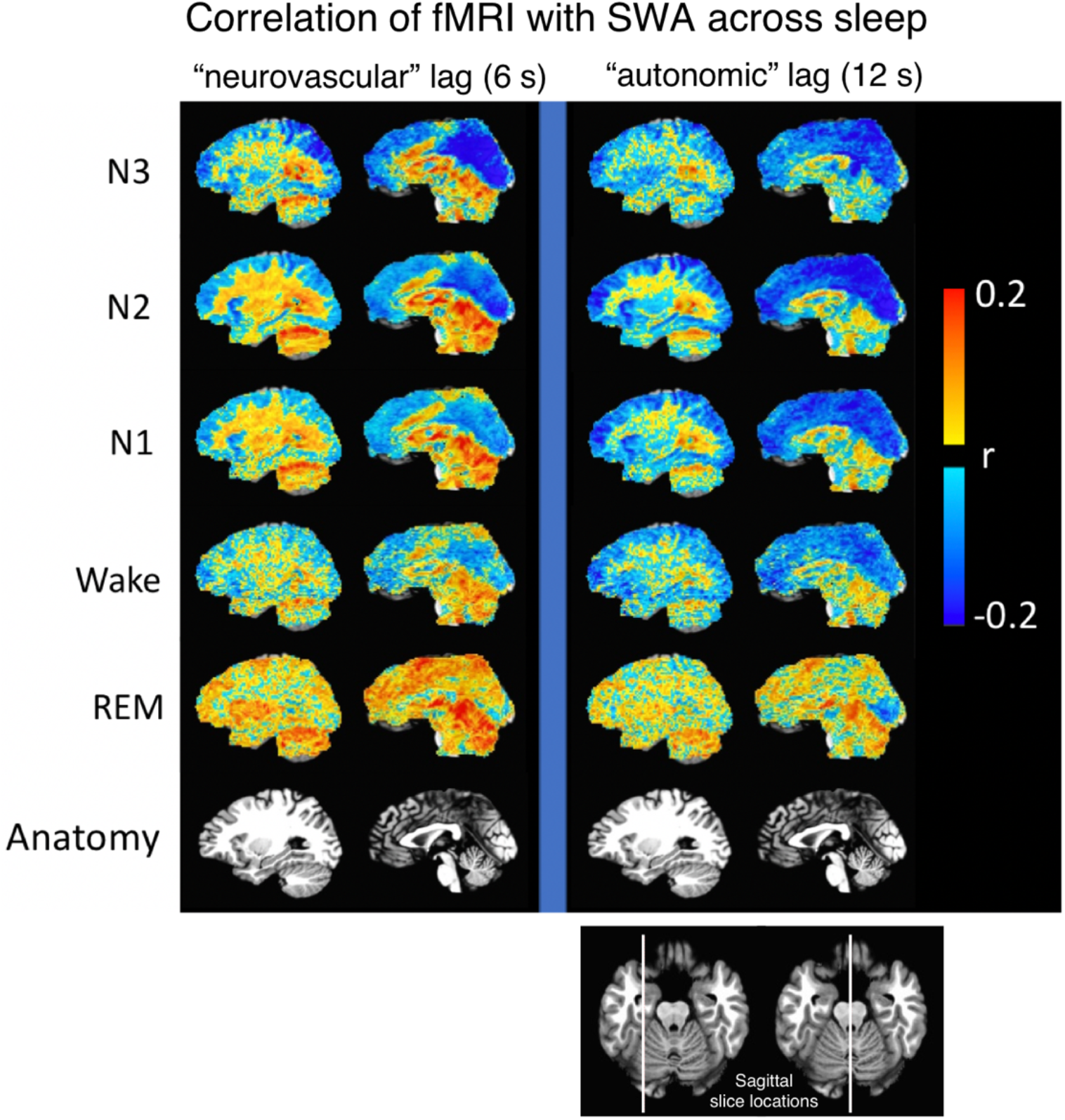
Spatially resolved correlation between slow wave activity (SWA) and fMRI signal across sleep stages. At a correlation lag of 6 s, where neuronal contributions to the fMRI signal are expected, relatively strong negative correlations are seen during N3, most prominently in posterior brain areas. Strong and more widespread correlations are also seen at a lag of 12 s, suggesting the SWA-fMRI association is mediated by an autonomic mechanism. This mechanism appears to be most relevant during N2 and N1.

**Fig. S2.**
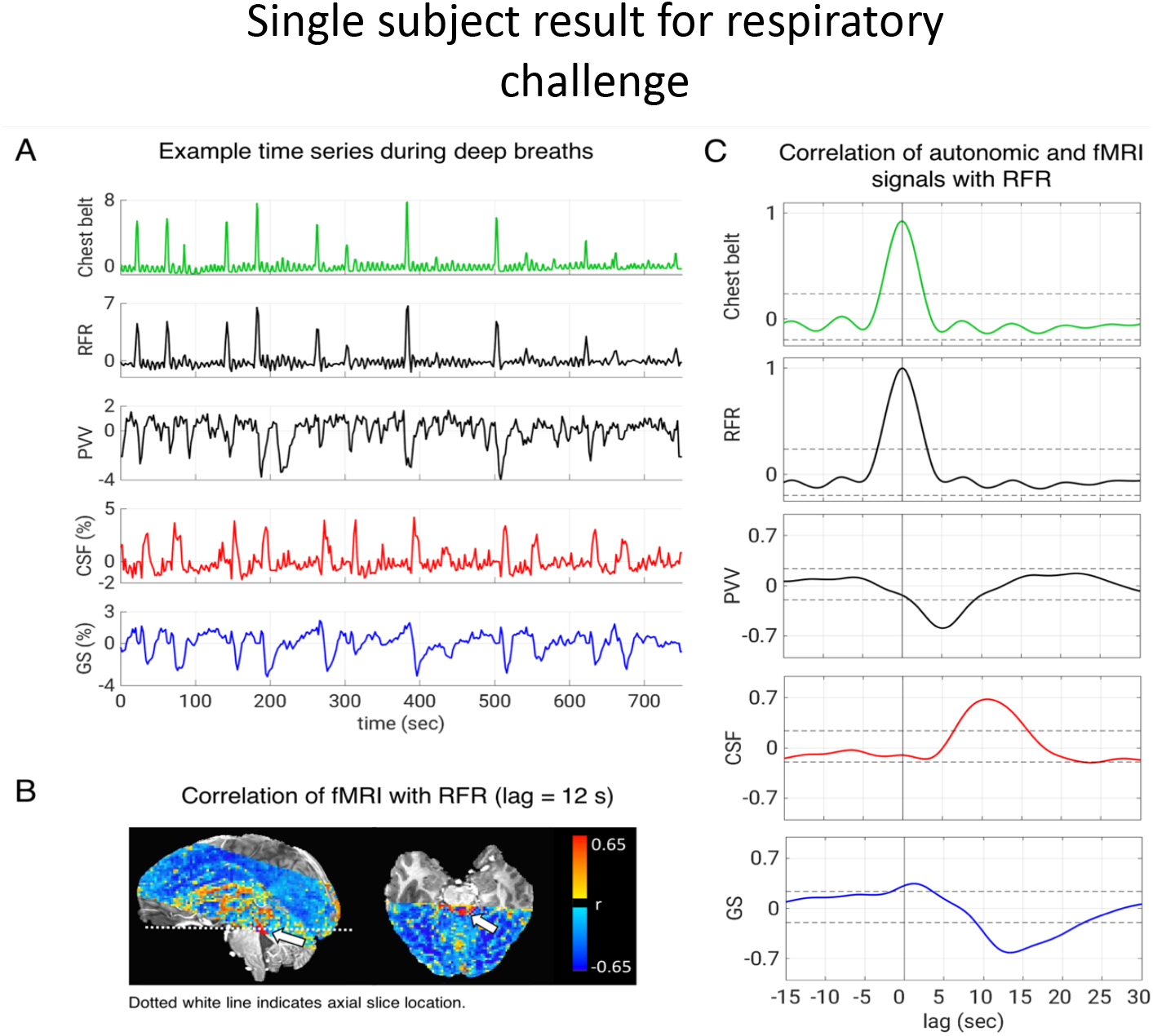
Single-subject example of the relationship between autonomic and fMRI signals during an inspiratory task. Similar to N1/N2 slow waves (K-complexes), periodic deep inspirations are associated with GS reductions and CSF inflow (**a,b**). Correlation of RFR with SWA (correlation lag=12 s) shows that SWA is associated with a global CBV decrease and an increase in CSF signal (arrow) in the cerebral aqueduct and the fourth ventricle (arrow) at the bottom of the fMRI slice stack (**b**). Delays between RFR and fMRI signals (**c**) are similar to those seen with K-complexes/SWA during sleep (see **Figs. 2-4**). Lags of significant correlations are indicated by horizontal line segments. GS = global signal, RFR = respiratory flow rate, PVV=peripheral vascular volume, CBV = cerebral blood volume. Autonomic signals in (**a**) are in units of temporal standard deviation. Horizontal dashed lines mark the significance threshold (p=0.05).

**Fig. S3.**
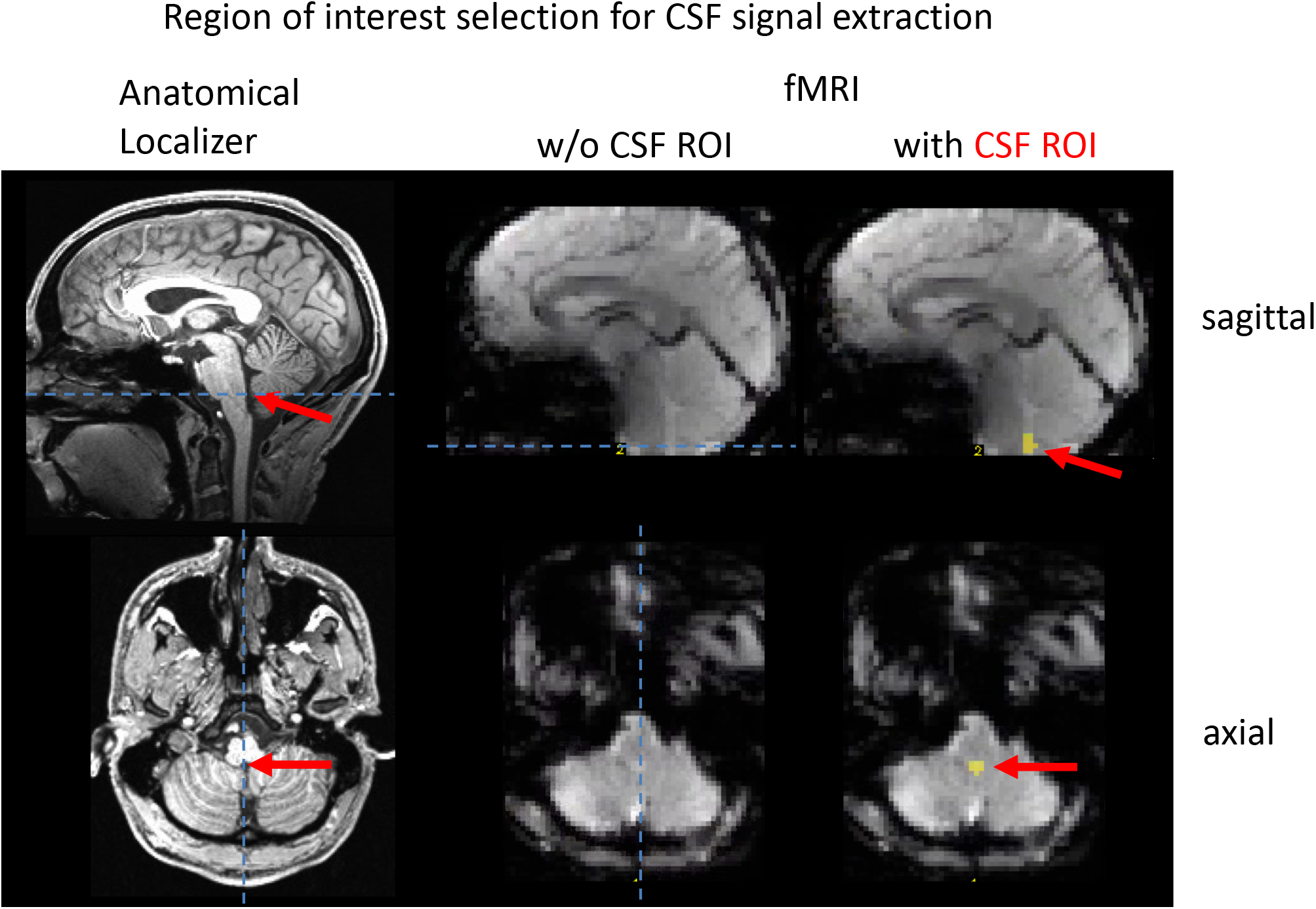
Selection of region of interest for fMRI CSF inflow signal shown on sagittal and sagittal MRI slices. The ROI is chosen in the CSF canal caudal to the fourth ventricle in three slices at the bottom of the slice stack most sensitive to inflow. Selection is based on the high signal intensity in the fMRI single time point images (left). Overlay of the selected ROI on fMRI images (right).

